# The F-box protein AFF1 regulates ARF protein accumulation to regulate auxin response

**DOI:** 10.1101/2021.04.25.441346

**Authors:** Hongwei Jing, David A. Korasick, Ryan J. Emenecker, Nicholas Morffy, Edward G. Wilkinson, Samantha K. Powers, Lucia C. Strader

**Author notes:** These authors contributed equally to this work (Hongwei Jing, David A. Korasick). Correspondence (L.C.S.).

## Abstract

Auxin critically regulates nearly every aspect of plant growth and development. Auxin-driven transcriptional responses are mediated through the AUXIN RESPONSE FACTOR (ARF) family of transcription factors. Although ARF protein stability is regulated via the 26S proteasome, molecular mechanisms underlying ARF stability and turnover are unknown. Here, we report the identification and functional characterization of an F-box E3 ubiquitin ligase, which we have named AUXIN RESPONSE FACTOR F-BOX1 (AFF1). AFF1 directly interacts with ARF19 and regulates its accumulation. Mutants defective in *AFF1* display ARF19 protein hyperaccumulation, attenuated auxin responsiveness, and developmental defects. Together, our data suggest a new mechanism, namely control of ARF protein stability, in regulating auxin response.

## INTRODUCTION

The plant hormone auxin plays a pivotal role in all aspects of the plant life cycle. The AUXIN RESPONSE FACTOR (ARF) family of transcription factors are central mediators of auxin transcriptional responses and several distinct mechanisms regulate ARF activity (*1*). The predominant regulator of ARF activity is by the interaction with, and repression by, Aux/IAAs. Under low auxin concentrations, Aux/IAAs directly repress ARF transcriptional activity. When auxin levels increase, auxin promotes complex formation of SCF^TIR1/AFB^ and Aux/IAAs; this coreceptor formation allows polyubiquitylation of Aux/IAAs, which are subsequently degraded through the 26S proteasome. Aux/IAA degradation relieves ARF protein repression, allowing ARF protein regulation of auxin-responsive genes. In addition to the repression-derepression paradigm, post-translational modifications (*2, 3*) and protein condensation (*4*) regulate ARF activity.

Accumulation of multiple ARFs is regulated by the 26S proteasome, including ARF1 (*5*), ARF2 (*6*), ARF6 (*7, 8*), ARF8 (*7*), and ARF17 (*7*). Despite these reports of ARF proteasomal regulation, the molecular mechanism regulating this process has yet to be identified and roles for ARF protein degradation were unknown.

To address this knowledge gap, we designed a forward genetics screen for mutants that display elevated YFP-ARF19 accumulation. We identified a mutant defective in a gene encoding an F-box protein, which we named *AUXIN RESPONSE FACTOR F-BOX1* (*AFF1*). AFF1 physically interacts with ARF19 and the closely-related ARF7. Further, *aff1* mutants hyperaccumulate ARF19 protein and display attenuated auxin responsiveness and morphological abnormalities. The increased ARF19 accumulation displayed by *aff1* results in increased ARF condensation, likely driving the observed attenuation of auxin responsiveness. Our results support a model in which the F-box protein AFF1 promotes ARF protein degradation to prevent inappropriate protein condensation to maintain auxin responsiveness.

## RESULTS

### ARF19 protein accumulation is proteasome-dependent

ARF7 and ARF19 are class-A ARFs that are likely transcriptional activators. These closely-related proteins coordinately play essential roles in auxin-mediated plant development (*9*). Similar to previous reports for ARF1 (*5*), ARF2 (*6*), ARF6 (*7, 8*), ARF8 (*7*), and ARF17 (*7*), we found that ARF7-HA and YFP-ARF19 protein accumulation increased upon application of the proteasome inhibitor MG132 (Fig. 1A), suggesting that ARF7 and ARF19 proteins are degraded in a proteasome-dependent manner. We further observed that YFP-ARF19 signal diminished over time, despite expression behind a constitutive promoter, when observed by microscopy (Fig. 1C, and fig. S1 and S2) or immunoblot analysis (Fig. 1, D and E, and fig. S1 and S2). Thus, ARF19 protein accumulation is developmentally regulated and dependent on the 26S-proteasome.

**Figure 1.**
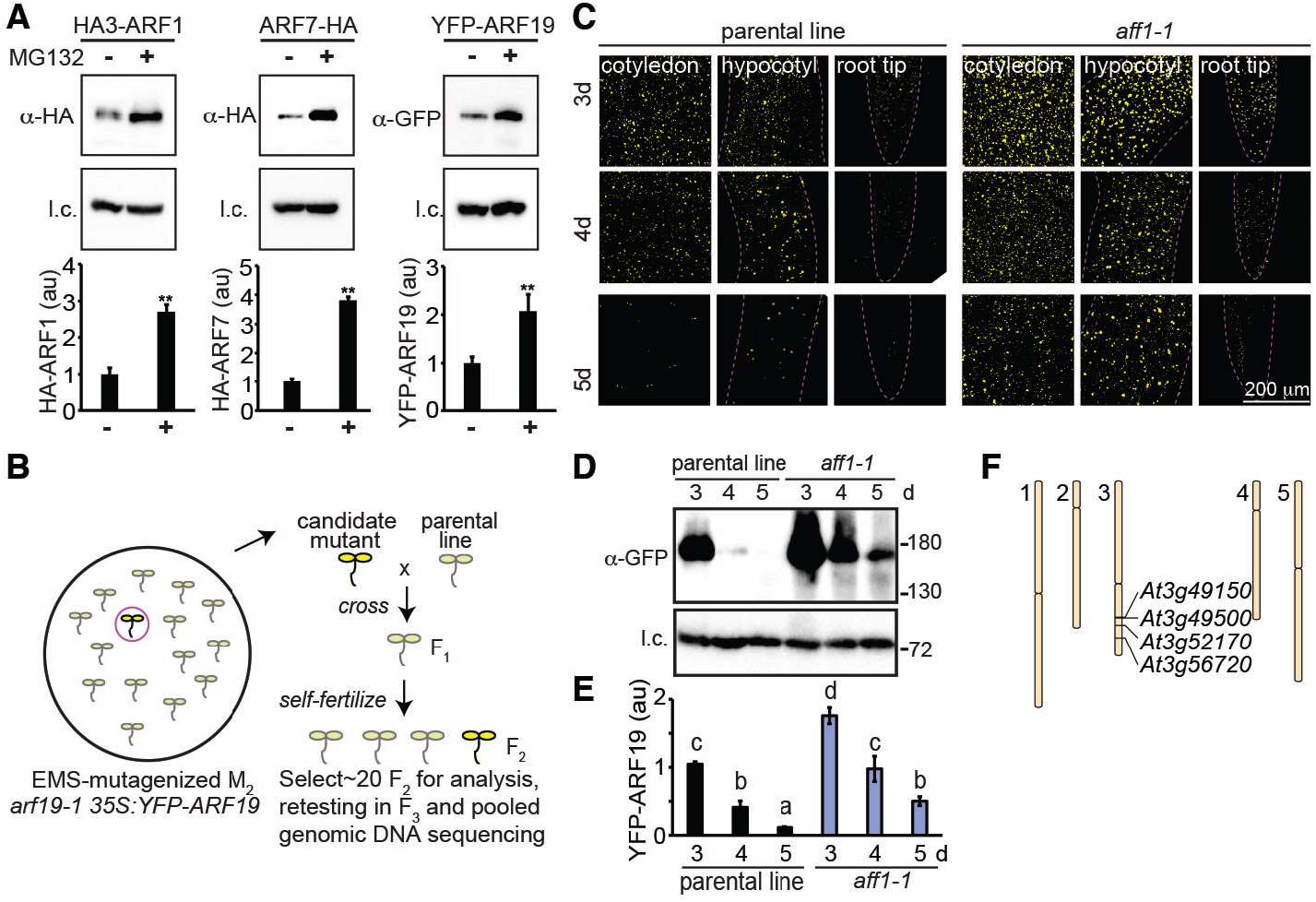
Identification of a YFP-ARF19 hyperaccumulation mutant. (A) Immunoblot analysis (image, top; quantification, bottom) of HA3-ARF1, ARF7-HA, and YFP-ARF19 treated with mock or MG132. Error bars = SD; \*\**P*<0.01 (Student’s *t*-test) (B) EMS-mutagenized M2 seed of *arf19-1 35S:YFP-ARF19* were screened for individuals with elevated YFP-ARF19 signal using a fluorescence dissecting microscope. Isolate DH8 was backcrossed to the parental line (P; *arf19-1 35S:YFP-ARF19*) and the resultant F2 individuals displaying YFP-ARF19 hyperaccumulation identified and used for whole-genome sequencing of bulk segregants (*10*). (C) Confocal microscopy images of YFP-ARF19 fluorescence from the parental line and *aff1-1* (DH8). Immunoblot analysis image (D) and quantification (E) of YFP-ARF19 protein levels in the parental line (P) and *aff1-1* (DH8). *P*<0.05 (LSD multiple range tests). (F) The four mutations identified in DH8 are on the lower arm of chromosome 3. Anti-HSC70 used for loading control (l.c.).

### Mutant screen and identification for YFP-ARF19 hyperaccumulators

To identify factors regulating ARF19 protein accumulation, we carried out a fluorescence-based screen of EMS-mutagenized *arf19-1 35S:YFP-ARF19* for individuals displaying elevated YFP-ARF19 signal (Fig. 1B). The isolate DH8 (*aff1-1*) displayed elevated YFP-ARF19 fluorescence (Fig. 1, C to E, and fig. S1 and S2). To identify the causative mutation, we used a whole genome sequencing of bulk segregants approach (Fig. 1B) (*10*), uncovering four homozygous, EMS-related mutations (Fig. 1F). Because ARF19 protein stability is regulated by the 26S proteasome (Fig.1A) and DH8 (*aff1-1*) hyperaccumulates ARF19 (Fig. 1, C to E, and fig. S1 and S2), we hypothesized that the mutation in *At3g49150*, encoding a putative F-box protein, was likely causative. We named this gene *AUXIN RESPONSE FACTOR F-BOX1* (*AFF1*) and our isolate *aff1-1* (Fig. 2A). The *aff1-1* mutant carries a C-to-T transition in the first exon of *AFF1*, resulting in the substitution of the Pro-93 with a Leu residue (Fig. 2A). The AFF1 protein contains an N-terminal F-box domain, a leucine rich repeat (LRR) region, and a C-terminal FBD motif (Fig. 2A).

**Figure 2.**
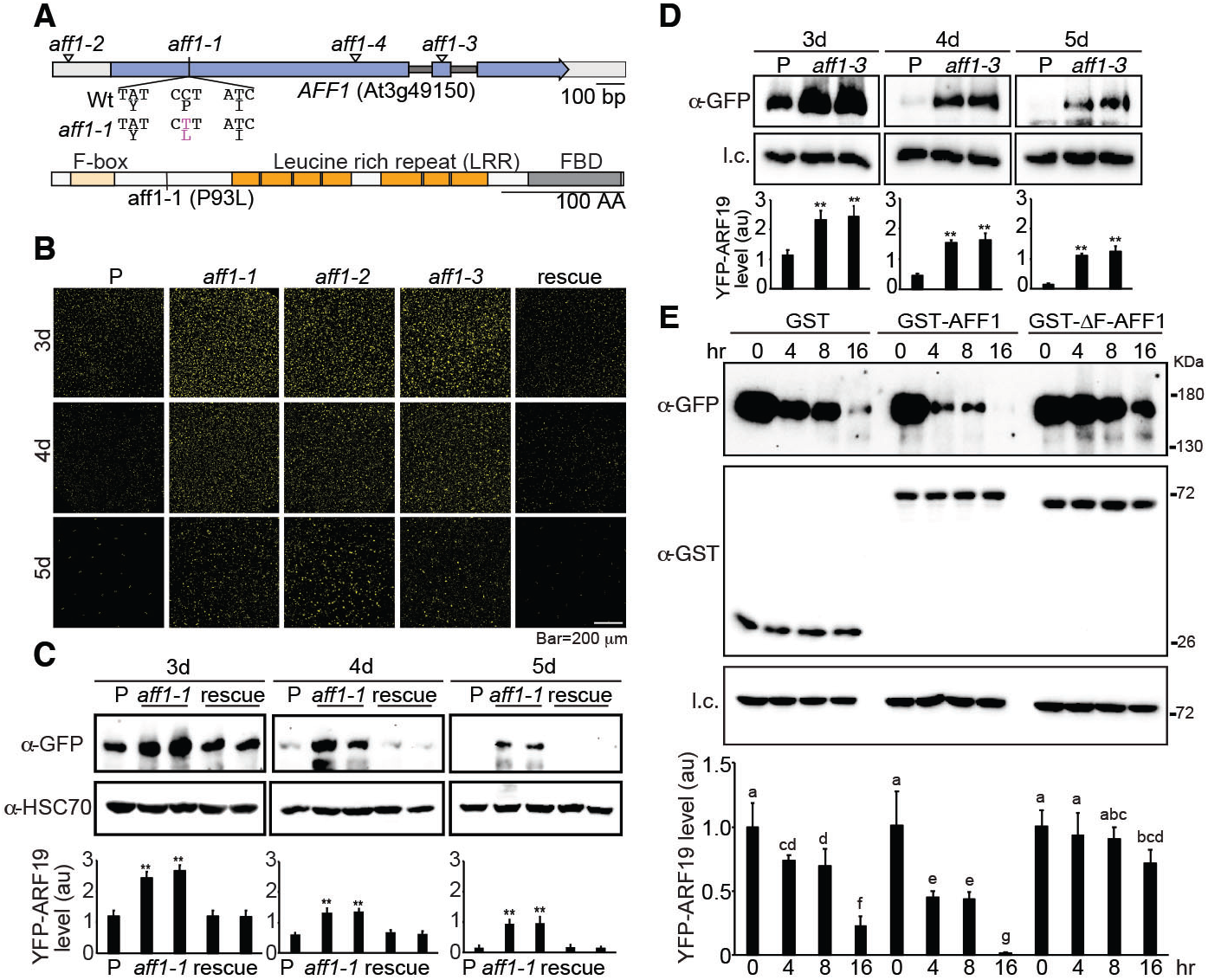
*At3g49150*/*AFF1* regulates YFP-ARF19 stability. (A) *At3g49150*/*AFF1* schematic depicting the exons (purple), UTRs (gray), and introns (black). Locations of the *aff1-1* point mutation mutant and *aff1-2* (Salk_053818), *aff1-3* (Salk_083453), and *aff1-4* (Sail_427_G06) insertion sites are indicated. *AFF1* encodes a putative F-box protein with an N-terminal F-box domain, leucine rich repeat (LRR) region, and C-terminal F-box domain (FBD) motif. (B) Confocal microscopy images of YFP-ARF19 fluorescence from *arf19-1 35S:YFP-ARF19* (P), *aff1-1*, *aff1-2*, and *aff1-3* and rescue line (*aff1-1 arf19-1 35S:YFP-ARF19 pAFF1:AFF1g*) cotyledons. (C, D) Immunoblot analysis of YFP-ARF19. (E) *In vitro* YFP-ARF19 degradation. Plant lysate from *aff1-1 arf19-1 35S:YFP-ARF19* were incubated with GST, GST-AFF1, or GST-ΔF-box-AFF1 recombinant proteins for the indicated times. Immunoblot analysis images (top) and quantification (bottom) of protein levels using the indicated antibodies. Anti-HSC70 used for loading control (l.c.). Error bars =SD; \*\**P*<0.01 (Student’s *t*-test). Letters indicate *P*<0.05 (LSD multiple range tests).

To verify whether the *At3g49150* mutation was causative in *aff1-1*, we identified three insertional alleles, which we named *aff1-2* (Salk_053818), *aff1-3* (Salk_083453), and *aff1-4* (Sail_427_G06) (Fig. 2A). YFP-ARF19 hyperaccumulates in these alleles, similar to the hyperaccumulation observed in *aff1-1* (Fig. 2, B to D, and Fig. S3). Moreover, we fully complemented *aff1-1* with a wild type copy of *AFF1*, confirming that the *At3g49150*/*AFF1* mutation is causative for the YFP-ARF19 hyperaccumulation observed in *aff1-1* (Fig. 2, B and C). *In vitro* YFP-ARF19 protein degradation assays showed that incubating plant lysate with GST-AFF1 recombinant protein increased YFP-ARF19 degradation compared to incubation with GST (Fig. 2E). Conversely, incubation of plant lysate with GST-ΔF-box-AFF1, a truncation of AFF1 that should be unable to incorporate into an SCF complex but retain the ability to interact with substrates, reduced YFP-ARF19 degradation (Fig. 2E), suggesting that this truncation protected ARF19 from endogenous degradation machinery. Thus, AFF1 regulates ARF19 protein accumulation.

### AFF1 interacts with ARF19 and ASK1 forming an SCF complex

Because AFF1 affects ARF19 accumulation, we explored whether AFF1 could directly interact with ARF proteins. We were unable to heterologously express full-length ARF19 protein; however, protein pull-down experiments revealed that GST-AFF1 and GST-ΔF-box-AFF1 interact with YFP-ARF19 protein from plant lysate (Fig. 3A). Further, YFP-ARF19 and ARF7-HA protein purified from plant lysate could interact with GST-AFF1 and GST-ΔF-box-AFF1 recombinant proteins, but not with GST (Fig. 3, B and C). Thus, AFF1 interacts with both ARF19 and its close homolog ARF7. In a bimolecular fluorescence complementation (BiFC) assay, we found that AFF1 interacted with ARF19 protein, but failed to interact with IAA7 (Fig. 3E, and fig. S4). Moreover, AFF1-ARF19 and ARF19-ARF19 interactions appeared to occur in the cytoplasmic ARF condensates whereas ARF19 and IAA7 appeared to interact primarily in the nucleus when transiently expressed in tobacco leaves (Fig. 3E, and fig. S4). Although the BiFC system artificially overexpresses proteins, these data are consistent with the possibility that AFF1 targets the cytoplasmic fraction of ARF proteins.

**Figure 3.**
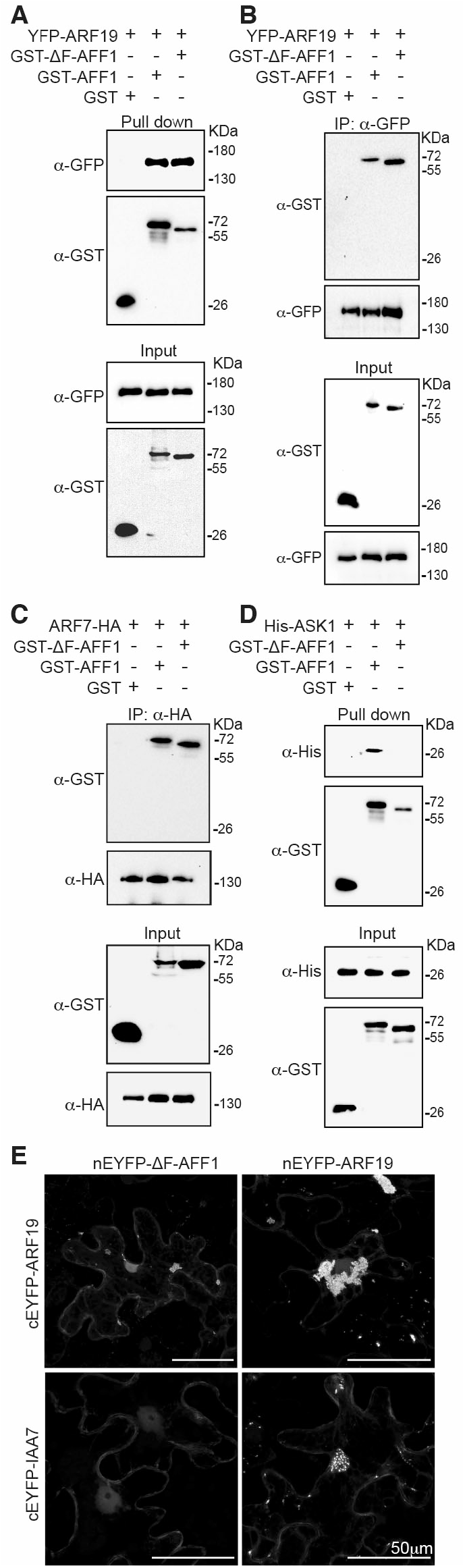
AFF1 interacts with ARF proteins and ASK1. (A) GST, GST-AFF1 or GST-ΔF-box-AFF1 recombinant proteins were incubated with *arf19-135S:YFP-ARF19* plant lysate. Pull-down fractions and inputs were examined by immunoblot analysis. (B) GST, GST-AFF1 or GST-ΔF-box-AFF1 were incubated with *arf19-1 35S:YFP-ARF19* plant lysate prior to immunoprecipitation with anti-GFP antibody. Immunoprecipitates and inputs were examined by immunoblot analysis. (C) GST, GST-AFF1 or GST-ΔF-box-AFF1 were incubated with *35S:HA-ARF7* plant lysate prior to immunoprecipitation with anti-HA antibody. Immunoprecipitates and inputs were examined by immunoblot analysis. (D) GST, GST-AFF1 or GST-ΔF-box-AFF1 were incubated with His-ASK1 prior to pull down. Pull-down fractions and inputs were examined by immunoblot analysis. (E) Bimolecular fluorescence complementation (BiFC; yellow) assay were used to analysis the protein interaction between nEYFP-ΔF-box-AFF1 and cEYFP-ARF19, nEYFP-ΔF-box-AFF1 and cEYFP-IAA7, nEYFP-ARF19 and cEYFP-ARF19, and nEYFP-ARF19 and cEYFP-IAA7. The nuclear marker WPP-mCherry (magenta) was co-expressed to determine nuclear signal. Extended dataset shown in Figure S4.

AFF1 is annotated as an F-box protein (Fig. 2A). F-box proteins typically contain an F-box domain, which allows incorporation into an SCF complex, and an additional domain that facilitates interaction with substrates. To test whether AFF1 can be incorporated in an SCF complex, in which F-box proteins must directly interact with ARABIDOPSIS SKP1-LIKE (ASK1), an adaptor connecting the subunit CULLIN 1 (CUL1) in the SCF complex (*11*), we examined AFF1 interactions with ASK1. In pull-down experiments, GST-AFF1, but not GST-ΔF-box-AFF1, interacted with heterologously-expressed His-ASK1 (Fig. 3D). Therefore, AFF1 can incorporate into an SCF E3 ubiquitin ligase complex. The direct interaction of AFF1 with ARF7 and ARF19 proteins, combined with our data that AFF1 regulates ARF19 accumulation, leads to a model in which SCF^AFF1^ regulates ARF19 protein stability.

### *AFF1* mutation exhibits developmental defects and altered auxin responsiveness

Similar to *ARF19* overexpression lines (*9*), morphometric analysis of *aff1* alleles revealed elongated and downward-curled leaves (Fig. 4A, and fig. S5), a phenotype often found in mutants defective in auxin signal transduction. In addition, *aff1* mutants displayed resistance to the inhibitory effects of the synthetic auxin 2,4-D on root elongation (Fig. 4, B and C). Consistent with these resistance phenotypes, *aff1* mutants displayed decreased auxin-responsive transcript accumulation in RNASeq-(Fig. 4, D to F) and Nanostring-(Fig. 4G) based analyses. These attenuated auxin response phenotypes were unexpected, as *aff1* accumulates elevated levels of the ARF19 protein which promotes auxin-responsive transcription, consistent with the possibility that the increased ARF19 protein found in *aff1* was not functional. Because ARF19 protein condensation was recently implicated in attenuating auxin responses (*4*), we examined ARF19 localization in the *aff1* mutant background, finding increased numbers of ARF19 condensates in the mutant background when compared to wild type (Fig 4H), suggesting that the attenuated auxin responses observed in *aff1* are caused by increased ARF19 condensation. Overall, our data support a model (Fig. 4I) in which the F-box protein AFF1 modulates ARF protein accumulation and thus condensation to regulate auxin responsiveness and plant growth and development.

**Figure 4.**
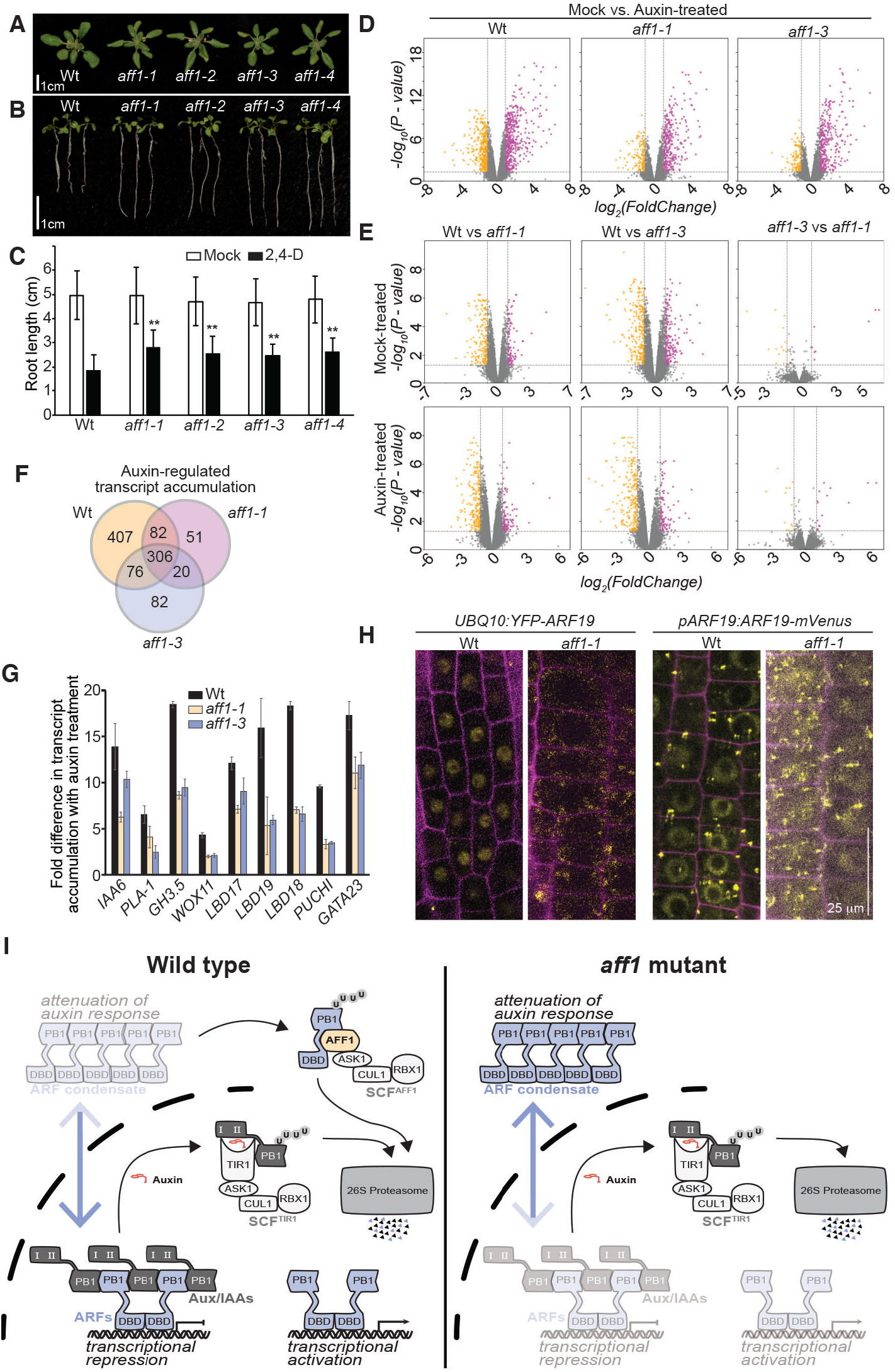
*aff1* exhibits developmental defects and attenuated auxin responsiveness. Photograph of 22d-old Wild type (Wt; Col-0), *aff1-1*, *aff1-2*, *aff1-3*, and *aff1-4* plants. (B) Photograph of 9d-old wild type, *aff1-1*, *aff1-2*, *aff1-3*, and *aff1-4* seedlings grown on media supplemented with 40 nM 2,4-D. (C) Mean (±SD; *n*=80) primary root lengths of 9d-old seedlings grown on media supplement with mock (EtOH) or 2,4-D. \*\**P*<0.01 (Student’s *t*-test).(D) Volcano plots displaying pairwise transcript accumulation differences after two hours of Mock (EtOH) or auxin (10 μM IAA) treatment in Wt (Col-0), *aff1-1*, and *aff1-3*. (E) Volcano plots displaying pairwise transcript accumulation differences between Wt (Col-0) and *aff1-1*, Wt and *aff1-3*, or *aff1-1* and *aff1-3* after 2-hour treatment with Mock (EtOH) or Auxin (10 μM IAA) for 2h. FDR ≤ 0.01. (F) Venn diagrams showing the number of genes that are overlap between the datasets of differentially expressed genes (FDR <0.01). (G) Relative transcript abundance (±SD, *n* = 3) of auxin response targets in Wt (Col-0), *aff1-1* and *aff1-3* with or without 10 μM IAA treatment for 2h. (H) Confocal images of Wt (Col-0) and *aff1-1* carrying *UBQ10:YFP-ARF19* or *pARF19:ARF19-mVenus* (yellow), with cell walls counterstained with propidium iodide (magenta). (I) A proposed model for the AFF1 role in regulating ARF stability. SCF^AFF1^ targets ARF19, and perhaps additional ARFs, to the proteasome. In the absence of AFF1, ARF protein hyperaccumulates, resulting in decreased nuclear ARF and thus attenuating auxin responses.

## DISCUSSION

The Arabidopsis genome contains over 700 genes encoding putative F-box proteins. SCF-mediated protein degradation plays critical roles in many aspects of plant growth and development (*12*). Here, we identify SCF^AFF1^ as a promoter of ARF19 destabilization to modulate auxin responses and plant development.

We found that elevated ARF19 levels counterintuitively resulted in attenuated auxin responses. Protein condensation and phase separation are concentration-dependent events (*13*). Further, ARF19 condensation attenuates auxin responsiveness (*4*). Our data raise the possibility that AFF1 acts as part of an ARF surveillance system to prevent inappropriately high ARF accumulation and condensation.

ARF transcription factors are divided into three ancient clades - termed Class-A, B, and C ARFs; Class-A ARFs are generally thought to be transcriptional activators whereas Class-B and C ARFs are generally thought to repress transcription (*14*). ARF7 and ARF19 are two class-A ARFs that are both regulated by the proteasome and directly interact with AFF1. Similarly, the class-A ARF5 and ARF8 also undergo proteasome-dependent degradation (*7*); whether this is through the activity of SCF^AFF1^ or through another mechanism remains unknown. Indeed, ARF19 is not fully stabilized in the *aff1* mutant backgrounds, suggesting that additional mechanisms regulate the stability of this transcription factor. Stability of the Class-B ARF1 is proteasome-dependent and its degradation rates are not altered in the *cul1* mutant background, suggesting that ARF1 proteasomal degradation via an alternative set of machinery (*5*). Thus, it seems likely that multiple mechanisms exist to regulate ARF protein accumulation.

We have not yet identified the ARF19 degron. However, ARF proteins which lack the PB1 domain (ARF17) (*7*), or are truncated to lack the DNA binding domain (ARF1) (*5*) are degraded in a proteasome-dependent manner. These findings raise the possibility that the ARF degron might lie within the intrinsically disordered middle region.

In summary, we have presented genetic and biochemical evidence demonstrating that the F-box protein AFF1 promotes ARF protein degradation to regulate auxin responses. Our uncovering this new mechanism that regulates ARF stability, ARF condensation, and auxin responses provides new insight into the mechanisms behind the complex web of auxin-regulated responses and opens new paths of investigation in auxin biology.

## AUTHOR CONTRIBUTIONS

## ACKNOWLEDGMENTS

We thank Joe Cammarata for critical comments on this manuscript and July Callis for providing HA3-ARF1. This research was funded by the USDA-NIFA Fellowship Program (MOW-2014-01877 to D.A.K.), the National Science Foundation Postdoctoral Research Program (IOS-1907098 to N.M.), and the National Institutes of Health (R35GM136338 to L.C.S.).

## METHODS

### Plant materials and phenotypic assays

All *Arabidopsis thaliana* lines were in the Columbia (Col-0) background, which was used as the wild type (Wt) in all experiments. For phenotypic assays, seeds were surface sterilized with 20% (v/v) bleach adding 0.01% (v/v) Triton X-100 for 15 min, then rinsed four times with sterile water (*15*). Sterilized seeds were suspended in 0.1% (w/v) agar and then stratified for 2 d at 4°C to promote uniform germination. After stratification, seeds were plated on plant nutrient (PN) media (*16*) solidified with 0.6% (w/v) agar and supplemented with 0.5% (w/v) sucrose (PNS) at 22 °C under continuous illumination. To analysis the leaf phenotypes in Wt and mutants, seeds were directly germinated in the soil. Images were taken after 22 d growth at 22 °C under continuous illumination. To examine root elongation in Wt and mutants, root lengths were measured from seedlings vertical-incubated media after 9 d of growth at 22 °C under continuous illumination.

### Vector construction and plant transformation

To create the parental line *arf19-1 35S:YFP-ARF19*, the coding sequence of ARF19 was amplified from cDNA using Pfx Platinum (Life Technologies). The PCR product was cloned into pENTR/D-TOPO (Life Technologies) to create pENTR-ARF19. The pENTR-ARF19 vector was recombined into the pEarleyGate104 (*35S:YFP-GW*) plasmid (*17*) using LR Clonase (Invitrogen). Recombinant plasmid was transformed into *Agrobacterium tumefaciens* strain GV3101 (*18*), and then transformed into *arf19-1* mutant plants via the floral dip method. Transformants were selected by T1 seeds and plating on plant nutrient media with sucrose (PNS) supplemented with 10μg/mL Basta. Subsequent generations were tested to identify lines homozygous for the transgene.

To create the rescue line *DH8 35S:AFF1genomic*, the genomic sequence of *AFF1* was cloned into pENTR/D-TOPO to create pENTR-AFF1g. The pENTR-AFF1g vector was recombined into the pMDC32 plasmid using LR Clonase. Recombinant plasmid was transformed into GV3101 and then used to transform into *DH8* mutant via the floral dip method. Transformants were selected by T1 seeds and plating on PNS supplemented with 25μg/mL hygromycin. Subsequent generations were tested to identify lines homozygous for the transgene.

The coding sequence of the AFF1 was synthesized and cloned into the pENTR/D-TOPO vector. The cDNA of AFF1 and △F-box-AFF1 were PCR amplified from the pENTR/D-TOPO vector, and then cloned into the *Bam*HI and *Sal*I sites of the pGEX4T-1 (Amersham Biosciences) to generate pGST-AFF1 and pGST-△F-box-AFF1 expression vectors. The coding sequence of ASK1 was PCR amplified from the Arabidopsis cDNA, and then cloned into the *Bam*HI and *Hind*III sites of the pET28a (Novagen) to make pHis-ASK1 expression vectors.

To create the bimolecular fluorescence complementation (BiFC) expression vectors, the pENTR-△F-box-AFF1, pENTR-TOPO-IAA7 and pENTR-ARF19 were recombined into the pSITE-nEYFP-C1 or pSITE-cEYFP-N1 (from ABRC) using LR Clonase. To create nuclear marker WPP-mCherry vector, the coding sequence of the WPP domain (amino acids 1-111) of the gene *RanGAP* (*At3g63130*) fused with mCherry was synthesized and cloned into the pENTR/D-TOPO vector. The ACT2 promoter was cloned into pENTR-WPP-mCherry using K*pn*I and X*ho*I restriction sites. Then the pENTR-ACT2-WPP-mCherry vector was recombined into the pMDC99 plasmid using LR Clonase to create pMDC99-ACT2-WPP-mCherry.

All primers used for plasmid construction are listed in Supplementary Table 1.

### EMS mutagenesis and mutant identification

To perform the mutant screen, nearly 5000 seeds of parental line *arf19-1 35S:YFP-ARF19* were ethyl methanesulfonate (EMS) mutagenized. The seeds were incubated with 0.24% (v/v) EMS for 16 hr, and then rinsed four times with sterile water. Mutagenized seeds were suspended in 0.1% (w/v) agar, then directly planted to soil in separate pots. M2 seeds from individual mutant pools were plated on PNS media growing for 8 d at 22°C under continuous illumination. Candidate mutants displaying elevated YFP-ARF19 signal using a fluorescence dissecting microscope were transferred to soil and allowed to self fertilize at 22°C under continuous illumination. Whole genome sequencing of bulk segregants approach was used to identify the causative mutation in DH8 mutant, which was described previously (*10*). We back-crossed DH8 three times into the wild type (Col-0) background. Nearly 500 progenies of a BC_3_F_3_ population were genotyped using PCR analysis to identify a single mutant *aff1-1*. The genotyping primers designed by dCAPS are listed in Supplementary Table 1.

### Confocal microscopy

For confocal images of plant lines, seedlings were mounted in water under a coverslip and imaged though a x40 lens on a Leica TCS SP8 confocal microscope.

### Immunoblot analysis

Immunoblot analysis was performed as described previously (*19*). Total cellular proteins were prepared by grinding plant materials in liquid nitrogen and then extracted in grinding buffer (50 mM Tris-HCl, pH 8.0, 150 mM NaCl, 1% (v/v) Nonidet P-40, 0.5% (w/v) sodium deoxycholate, 0.1% (w/v) SDS, 1 mM phenylmethylsulfonyl fluoride and 1% (v/v) protease inhibitors cocktail (Sigma-Aldrich, P9599). After heating at 100°C for 10 min, the samples were then subjected to SDS-polyacrylamide gel electrophoresis. After the run, proteins were transferred onto a nitrocellulose membrane, and then detected with 1:5000 of the indicated primary antibody. The blot was incubated with a secondary antibody (goat anti-mouse IgG-HRP or goat anti-rabbit IgG-HRP, Santa Cruz Biotechnology) at 1:5000 dilution. The signal was detected using a WesternBright ECL HRP substrate kit (Advansta) according to the manufacturer’s instructions. Arabidopsis HSC70 was used as a loading control. The target bands and loading control bands were quantified using ImageJ and the mean values of 3-5 independent experiments were presented with statistical analysis (LSD multiple range tests or Student’s *t*-test) of significant differences when applicable.

For MG132 (Sigma-Aldrich, M8699) treatment, *UBQ10:HA_3_-ARF1*, *35S:ARF7-HA*, or *arf19-1 35S:YFP-ARF19* were grown on PNS media for 3 d at 22 °C under continuous illumination. Seedlings were then transferred to liquid PN supplemented with either DMSO (Mock) or 50 μM MG132 for 16h. Then the samples were collected for following immunoblot analysis. Three independent experiments were used for quantitative analysis.

### Bimolecular fluorescence complementation (BiFC) assay

Bimolecular fluorescence complementation (BiFC) experiments were conducted as previously described (*20*). Briefly, the resulting binary expression vectors were transformed into *Agrobacterium* strain GV3101. Collected cells were washed and resuspended to OD_600_ of approximately 1.0 with the infiltration solution (10 mM MES (pH 5.6), 10 mM MgCl_2_, and 1 mM acetosyringone). *Agrobacterial* cells carrying various expression vectors with the p19 strain were co-infiltrated into 3-week-old *Nicotiana benthamiana* leaves. Empty vectors were used as negative controls. After the infiltration, plants were placed at 22 °C for 3 d and the YFP and mCherry fluorescent signals were detected using Leica TCS SP8 confocal microscope. The experiment was repeated three times with independent biological replicates.

### Pull-down assay

Protein pull-down assay was performed as described with minor modifications (*21*). To analysis the interaction between ARF19 or ARF7 with AFF1, plant samples from *arf19-1 35S:YFP-ARF19* or *35S:ARF7-HA* were ground in liquid nitrogen, and then extracted in grinding buffer (50 mM Tris-HCl, pH 7.5, 150 mM NaCl, 10 mM MgCl_2_, 10% (v/v) glycerol, 0.1% (w/v) Nonidet P-40, 1 mM phenylmethylsulfonyl fluoride, 1% (v/v) protease inhibitors cocktail (Sigma-Aldrich, P9599) and 10 μM MG132. Purified GST, GST-AFF1, and GST-△F-box-AFF1 proteins were immobilized on GST beads (Glutathione Agarose; ThermoFisher). Immobilized agarose beads containing 2 μg GST, GST-AFF1, or GST-△F-box-AFF1 fusion proteins were mixed with nearly 1.0-2.0 mg total cellular proteins from *arf19-1 35S:YFP-ARF19* or *35S:ARF7-HA* at 4 °C for 2 hr. The beads were collected by centrifugation and then washed six times with washing buffer (10 mM phosphate buffer saline, pH 7.4, 150 mM NaCl, 0.2% (v/v) Triton X-100 and 1 mM phenylmethanesulfonyl fluoride) at 4 °C. The beads were resuspended in SDS-polyacrylamide gel electrophoresis sample buffer and then analyzed by immunoblot.

Using the Pull-down assay to detect the interaction between GST-AFF1 and His-ASK1, immobilized agarose beads containing 2 μg GST, GST-AFF1, or GST-△F-box-AFF1 fusion proteins were mixed with 2 μg His-ASK1 at 4 °C for 2 hr. The beads were collected after washing six times to do the SDS-polyacrylamide gel electrophoresis, and then analyzed by immunoblot.

### Co-immunoprecipitation assay

The Co-IP experiments were performed according to previously described methods with minor modifications (*22*). To prepare total cellular proteins, plant samples were grinded in liquid nitrogen, and then extracted in grinding buffer (50 mM Tris-HCl, pH 7.5, 150 mM NaCl, 10 mM MgCl_2_, 10% (v/v) glycerol, 0.1% (v/v) Nonidet P-40, 1 mM phenylmethylsulfonyl fluoride, 1% (v/v) protease inhibitors cocktail (Sigma-Aldrich, P9599) and 10 μM MG132. The extracts containing 1.0-2.0 mg total cellular proteins were incubated with 10 μl anti-GFP or anti-HA antibodies for 1 hr at 4 °C with gentle shaking. After that, the Dynabeads Protein G (50 μl, ThermoFisher) were added and mixed with 2 μg GST, GST-AFF1, or GST-△F-box-AFF1 fusion proteins for an additional 2 hr at 4 °C. The immunoprecipitates were washed six times with 1 ml washing buffer (grinding buffer without MG132) and then used for immunoblot.

### *In vitro* turnover assay

The analysis of YFP-ARF19 protein degradation *in vitro* was performed as described methods with minor modifications (*19*). In brief, total protein extracts were prepared from 3 d-old parental line *arf19-1 35S:YFP-ARF19* grown in PNS medium using ice-cold extraction buffer (50 mM Tris-HCl, pH 7.5, 150 mM NaCl, 0.01% (v/v) Triton X-100 and 1 mM phenylmethanesulfonyl fluoride). The crude extracts (1 mg proteins) were mixed with 2 μg of purified GST, GST-AFF1, or GST-△F-box-AFF1 recombinant proteins in a total volume of 600 ml. The mixture was incubated at 4 °C with gentle agitation and 100 ml of each sample was collected at the indicated time points and then analysed by immunoblotting.

### Quantitaive reverse transcription-PCR (qRT-PCR)

Total RNA was prepared using the RNeasy Plant Mini Kit (Qiagen). Quantitative reverse transcription-PCR (qRT-PCR) was performed using the iTaq™ Universal SYBR^®^ Green Supermix (Bio-Rad) according to the manufacturer’s instructions. The reactions were run in a CFX96 REAL-Time PCR Detection System (Bio-Rad). The relative expression level of the target genes was analysed with the delta-delta Ct method and normalized to the expression level of *ACT7*. All of the experiments were repeated for at least twice (two biological repeats with three technical repeats for each experiment). The primers used for qRT-PCR are listed in Supplementary Table 1.

### RNA-Seq experiment

RNA-Seq experiment were performed according previously described methods (*4*). Col-0 (Wt), *aff1-1*, and *aff1-3* (Salk_083453) were grown on PNS media for 4 d at 22 °C under continuous illumination. Seedlings were then transferred to liquid PN supplemented with either ethanol (Mock) or 10 μM IAA for 2h. Three repeated treatments were carried out for each line. Total RNA was isolated using the RNeasy Plant Mini Kit (Qiagen). RNA samples were then sequenced using the Epicentre Ribo-Zero Gold system according to manufacturer’s protocol, indexed, pooled, and sequenced across three 1×50bp lanes on a single flow-cell on an Illumina HiSeq 3000. RNA-seq reads were demultiplexed and aligned to the Ensembl release 23 (TAIR 10) top-level assembly with STAR version 2.0.4b. Gene counts were derived from the number of uniquely aligned unambiguous reads by Subread:featureCount version 1.4.5. Sequencing performance was assessed for total number of aligned reads, total number of uniquely aligned reads, and genes detected.

All gene counts were imported into the R/Bioconductor package EdgeR and TMM normalization size factors were calculated to adjust for samples for differences in library size. Ribosomal genes and genes not expressed in any sample greater than one count-per-million were excluded from further analysis. In addition, genes not expressed in at least 2 out of the 3 samples were not considered for downstream analysis. The TMM size factors and the matrix of counts were then imported into R/Bioconductor package Limma. Performance of the samples was assessed with a Spearman correlation matrix and Multi-Dimensional Scaling plot (S6a,b). Weighted likelihoods based on the observed mean-variance relationship of every gene and sample were then calculated for all samples with the voomWithQualityWeights function and gene performance was assessed with plots of residual standard deviation of every gene to their average log-count with a robustly fitted trend line of the residuals (S6c). A generalized linear model was then created to test for gene level differential expression and the results were filtered for only those genes with Benjamini-Hochberg false-discovery rate adjusted p-values less than or equal to 0.05.

For volcano plots and heat maps, data was imported using the Pandas python package. For volcano plots, the bioinfokit Python package was used (visuz.gene_exp.volcano), and vertical lines represent the LFC of 1.5 and the horizontal lines represent adjusted p-values of 0.05. For heat maps, the Python package Seaborn was used (seaborn.clustermap) with a custom color map using matplotlib.colors.

### NanoStrings Analysis

NanoStrings analysis experiment were performed according previously described methods (*4*). Col-0 (Wt), *aff1-1*, and *aff1-3* (Salk_083453) were grown on PNS media for 4 d at 22 °C under continuous illumination. Seedlings were then transferred to liquid PN supplemented with either ethanol (Mock) or 10 μM IAA for 2h. Three repeated treatments were carried out for each line. Total RNA was isolated using the RNeasy Plant Mini Kit (Qiagen). NanoString nCounter analysis was performed using 80 ng total RNA and carried out using the nCounter Digital Analyzer (NanoStrings Technologies; Seattle, WA) at the McDonnell Genome Institute at Washington University in St. Louis. In addition to 8 negative-control and 6 positive-control probes, two genes *TUB4* (*At5g44340*) and *PP2C* (At1g13320) were used as references for normalization. Data was analyzed using the nSolver Analysis software.

---

**Figure S1.**
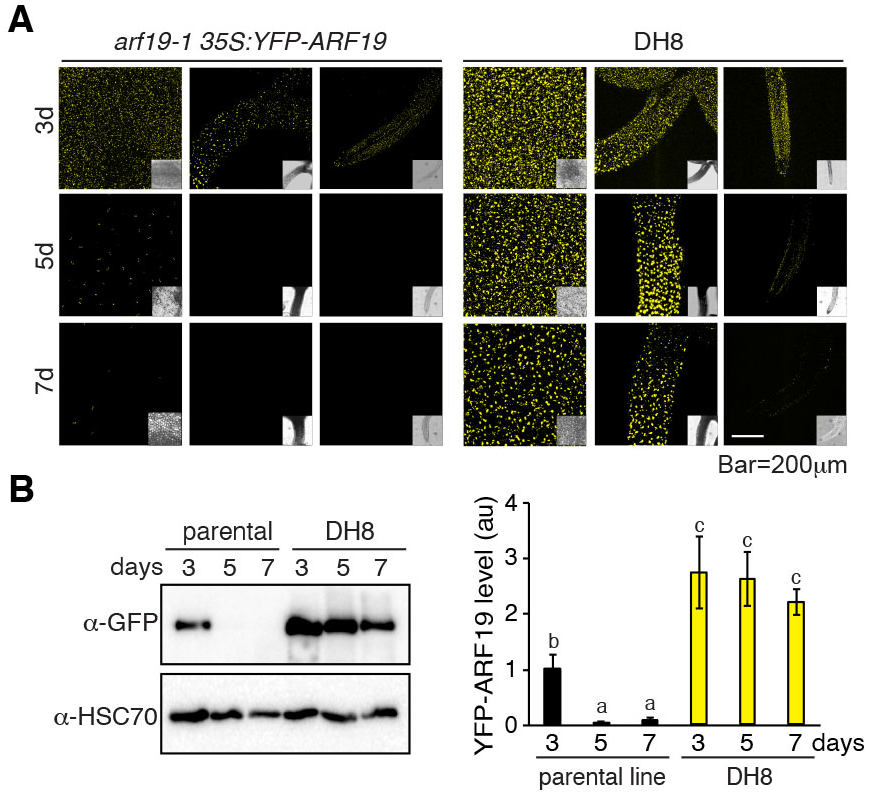
DH8 hyperaccumulation of YFP-ARF19. (A) Confocal microscopy images of YFP-ARF19 fluorescence from parental line (*arf19-1 35S:YFP-ARF19*) and DH8 mutant in cotyledon, hypocotyl, and root tip tissues at 3 days, 5 days, and 7 days. Scale bar, 200 μm. (B) Immunoblot analysis the YFP-ARF19 protein levels same as the panel (A) over a time course. (C) Quantitative analysis of the relative levels of YFP-ARF19 proteins same as the panel (B) (the average values obtained from three independent experiments). Loading control (l.c.) is verified by the analysis of HSC70 protein. Error bars = SD.; different letters indicate *P*<0.05 (LSD multiple range tests).

**Figure S2.**
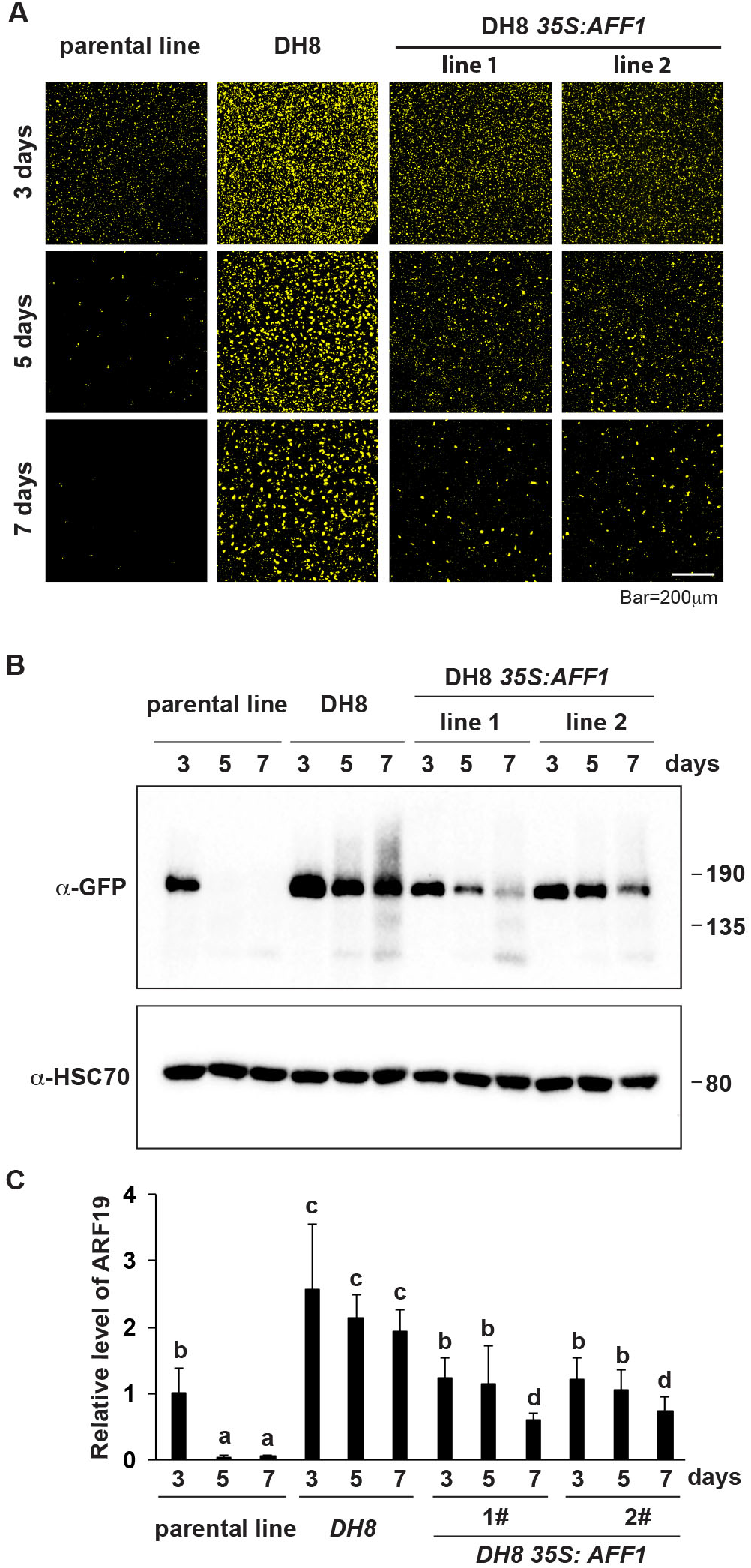
AFF1 complement the DH8 hyperaccumulation of YFP-ARF19 phenotype. (A) Confocal microscopy images of YFP-ARF19 fluorescence from parental line, DH8 and DH8 *35S:AFF1* in cotyledon at 3 days, 5 days, and 7 days. Scale bar, 200 μm. (B) Immunoblot analysis the YFP-ARF19 protein levels same as the panel (A) over a time course. Loading control (l.c.) is verified by the analysis of HSC70 protein. (C) Quantitative analysis of the relative levels of YFP-ARF19 proteins is presented below the blots (the average values obtained from three independent experiments). Error bars = SD; different letters indicate *P*<0.05 (LSD multiple range tests).

**Figure S3.**
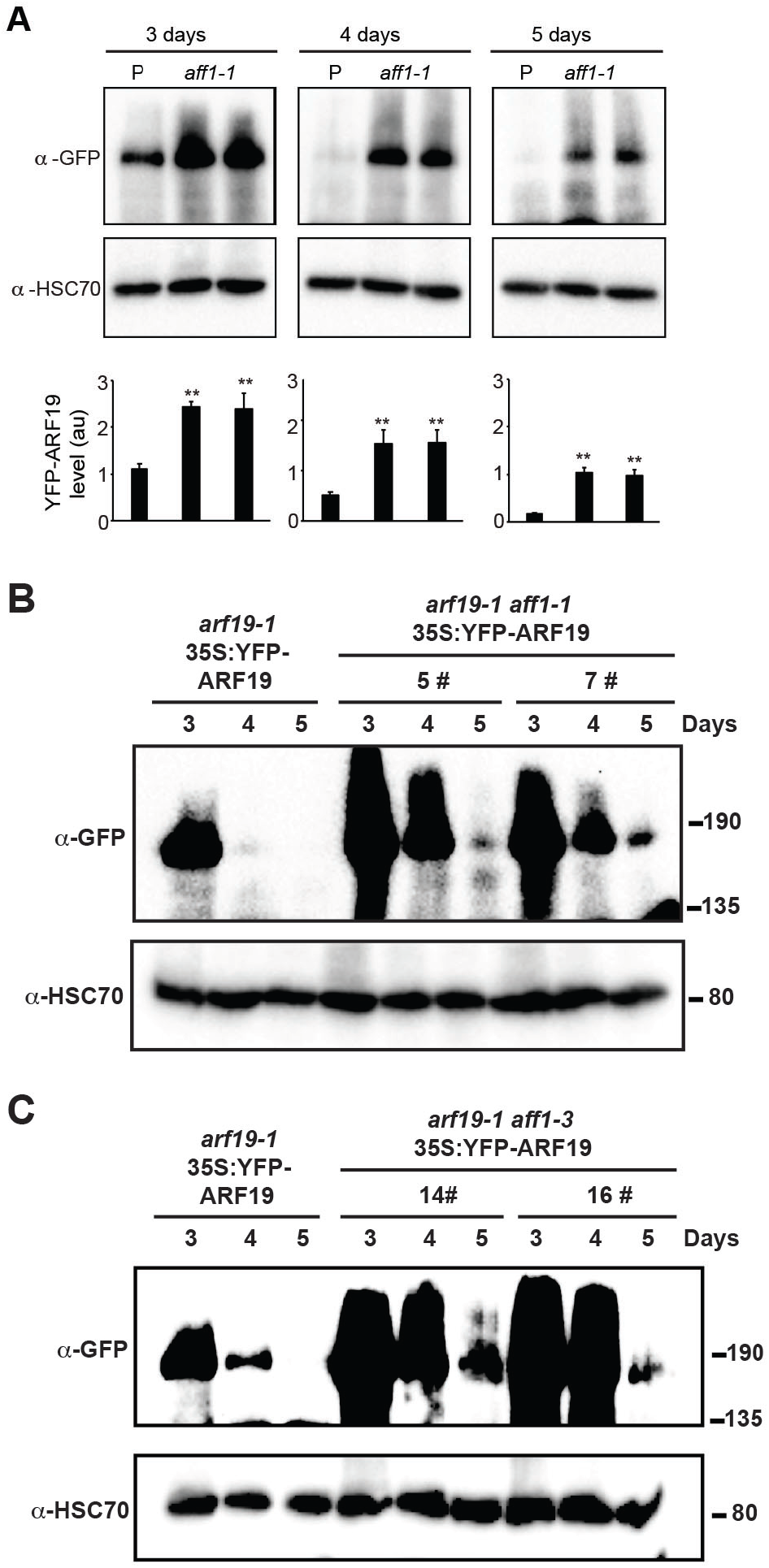
Hyperaccumulation of YFP-ARF19 in *aff1* mutants. (A) Immunoblot analysis the YFP-ARF19 protein levels in the parental line and the mutant *aff1-1* over a time course. Loading control (l.c.) is verified by the analysis of HSC70 protein. Quantitative analysis of the relative levels of YFP-ARF19 proteins is presented below the blots (the average values obtained from three independent experiments). Error bars indicate s.d.; \*\**P*<0.01 (Student’s *t*-test). (B, C) Immunoblot analysis the YFP-ARF19 protein levels in the parental line and the mutant *aff1-1* and *aff1-3* over a time course. Loading control (l.c.) is verified by the analysis of HSC70 protein.

**Figure S4.**
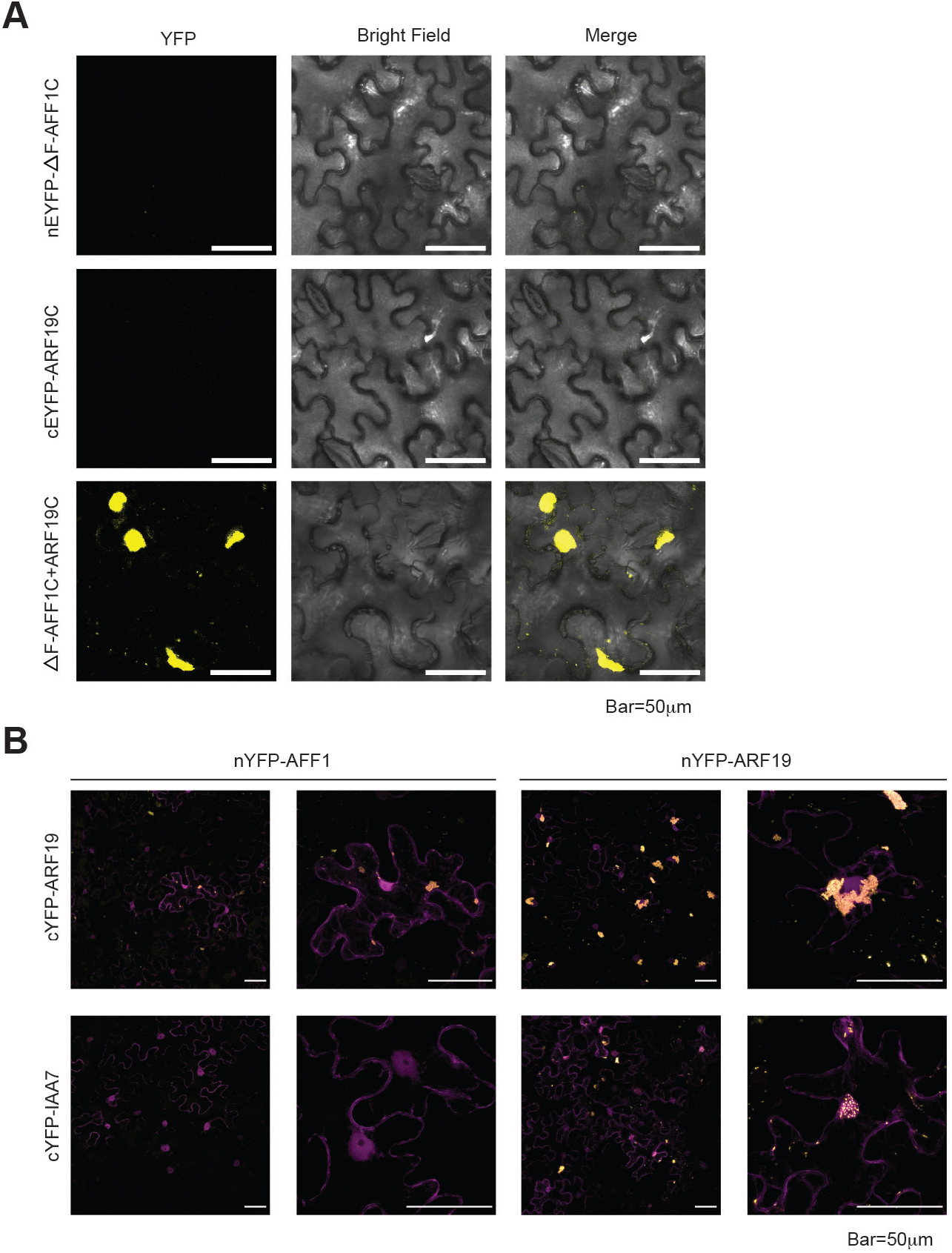
Interaction of ARF19 with ΔF-box-AFF1 using BiFC experiment. (A) BiFC assays were used to determine the interaction between ARF19 with ΔF-box-AFF1 when transiently expressed in tobacco leaves. nEYFP-ΔF-box-AFF1C denotes expression of the EYFP N-terminal fusion with ΔF-box-AFF1C construct. cEYFP-ARF19C denotes expression of the EYFP C-terminal fusion with ARF19C construct. Scale bar, 50 μm (B) BiFC assay were used to analysis the interaction between nEYFP-△F-box-AFF1 and cEYFP-ARF19, between nEYFP-ΔF-box-AFF1 and cEYFP-IAA7, between nEYFP-ARF19 and cEYFP-ARF19, and between nEYFP-ARF19 and cEYFP-IAA7. Nuclear marker WPP-mCherry was co-expressed in the tobacco leaves. Scale bar, 50 μm. Right panel images of each treatment were same as the Fig. 3E.

**Figure S5.**
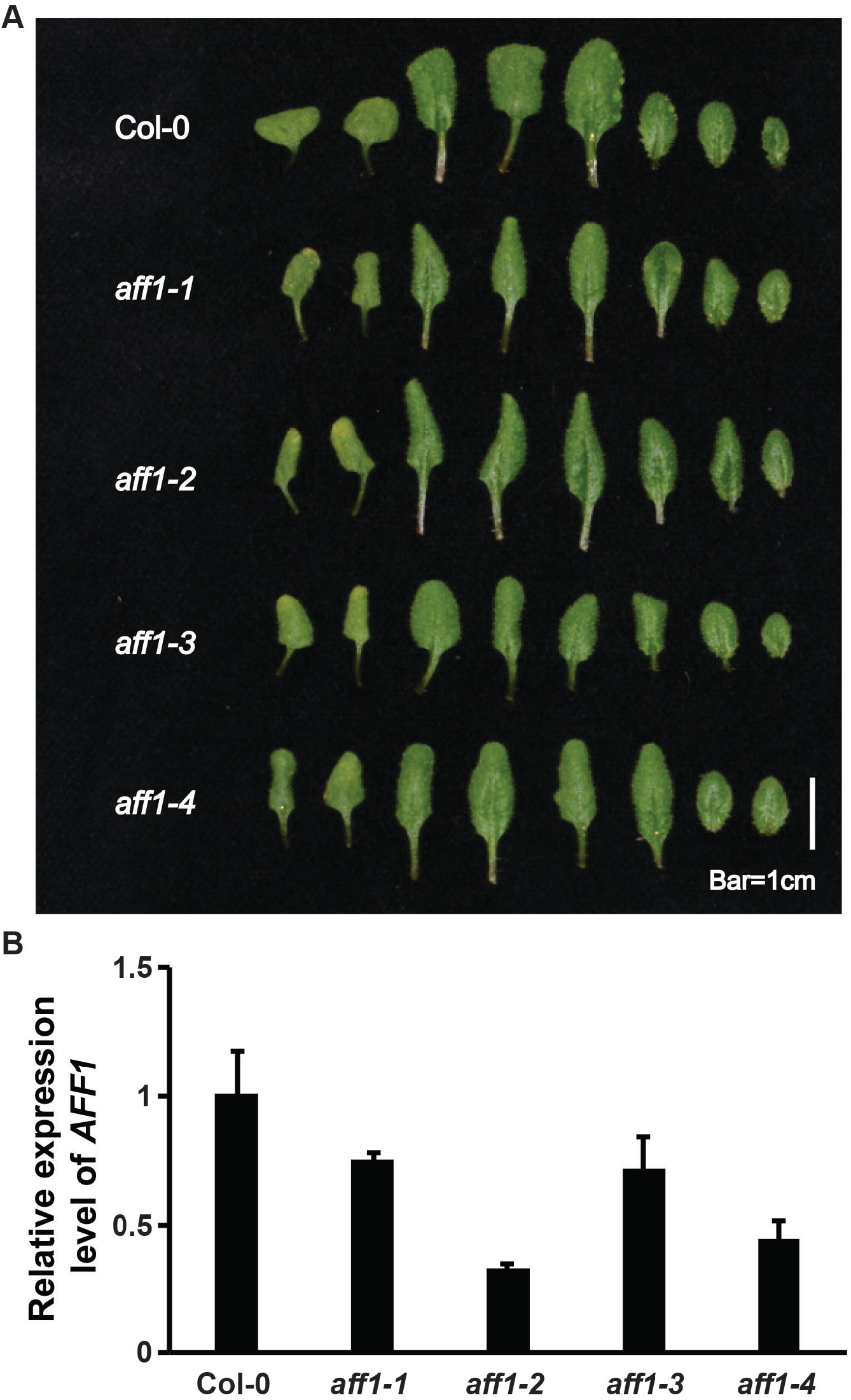
*AFF1* mutation exhibits developmental defects. (A) Leaves phenotypes of 22d-old wild type (Col-0), *aff1-1*, *aff1-2*, *aff1-3*, and *aff1-4* plants grown in soil. Scale bar, 1 cm. (B) Analysis of the expression level of *AFF1* in wild type, *aff1-1*, *aff1-2*, *aff1-3*, and *aff1-4* by qRT-PCR. Error bars indicate s.d.

**Figure S6.**
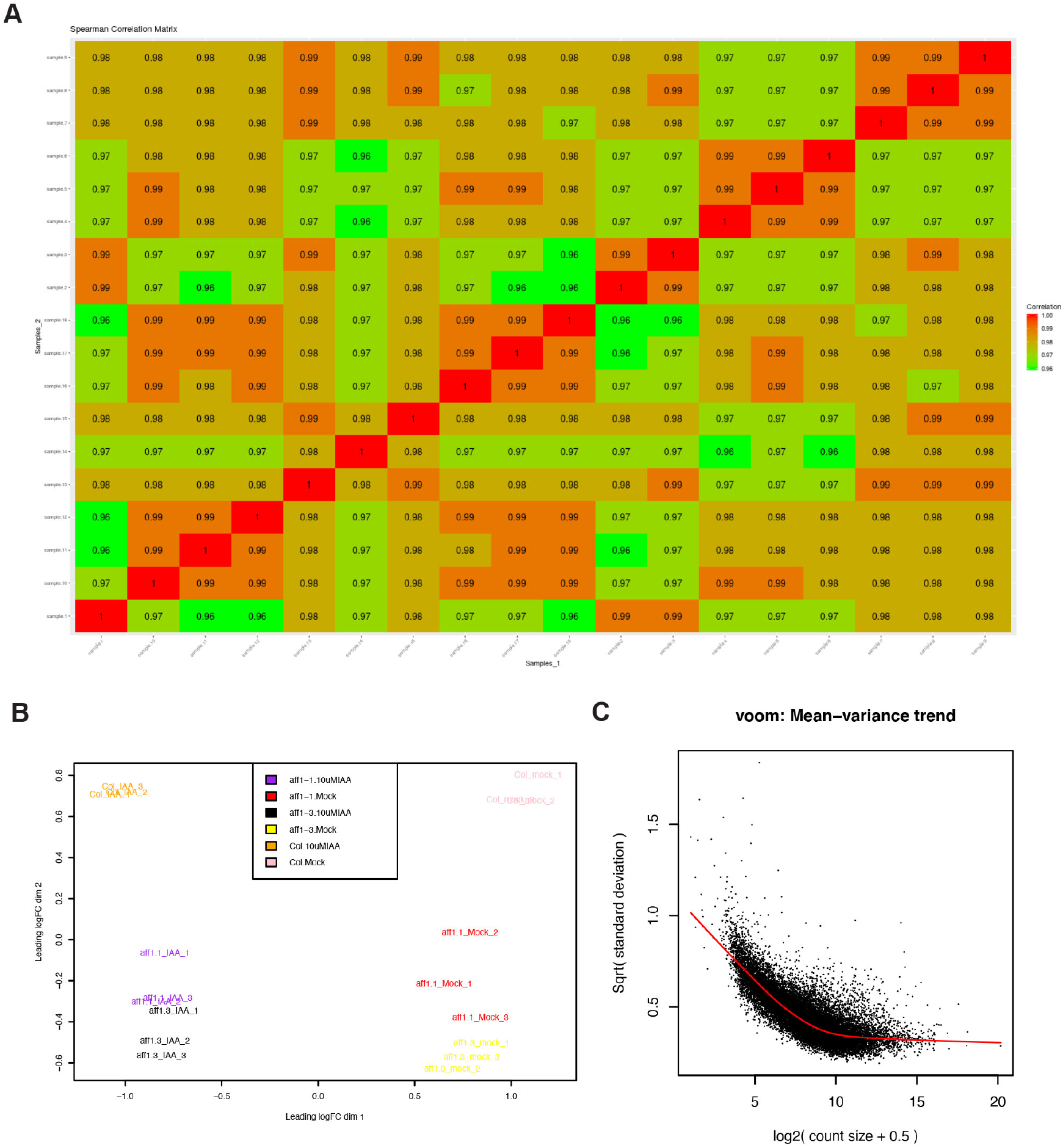
RNA-seq sample quality. (A) Matrix of Spearman correlations of all detected genes greater than 1 count-per-million in at least 3 samples relative to each other. Samples of the same condition have very high correlations as expected across the diagonal with no outliers. (B) The quality of the samples in a multi-dimensional scaling plot of the leading log fold- changes. Samples for each condition cluster very tightly with each other with good cluster separation between samples of different conditions based on expression profiles. (C) Scatter plot of the empirically derived fitted and trended mean-variance relationship across all genes.

